# Signatures of co-evolution and co-regulation in the CYP3A and CYP4F genes in humans

**DOI:** 10.1101/2023.02.23.529697

**Authors:** Alex Richard-St-Hilaire, Isabel Gamache, Justin Pelletier, Jean-Christophe Grenier, Raphael Poujol, Julie G Hussin

## Abstract

Cytochromes P450 (CYP450) are hemoproteins generally involved in the detoxification of the body of xenobiotic molecules. They participate in the metabolism of many drugs and genetic polymorphisms in humans have been found to impact drugs responses and metabolic functions. In this study, we investigate the genetic diversity for *CYP450* genes. We found that two clusters, *CYP3A* and *CYP4F*, are notably differentiated across human populations with evidence for selective pressures acting on both clusters: we found signals of recent positive selection in *CYP3A* and *CYP4F* genes and signals of balancing selection in *CYP4F* genes. Furthermore, unusual linkage disequilibrium pattern is detected in both clusters, suggesting co-evolution of genes within clusters. Several of these selective signals co-localize with expression quantitative trait loci, which suggest co-regulation and epistasis within these highly important gene families. We also found that SNPs under selection in Africans within the *CYP3A* cluster are associated to *CYP3A5* expression levels which are causally associated with reticulocytes count, as established by mendelian randomization. Furthermore, as the *CYP3A* and *CYP4F* subfamilies are involved in the metabolism of nutrients and drugs, our findings linking natural selection and gene expression in these gene clusters are of importance in understanding population differences in human health.

## 2 Introduction

In the last decades, it has become clear that every individual has their own “fingerprint” of alleles encoding drug-metabolizing enzymes, playing central roles in the metabolism of endogenous and exogenous compounds. Early in the 1960s, it was established that hydrophobic molecules are first modified by oxidation and subsequently excreted as water-soluble forms, two distinct steps now described as phases I and II. Phase I is performed mainly by Cytochromes P450 (CYP450) enzymes, able to catalyze a considerable variety of oxidation reactions for many structural classes of chemicals (including the majority of drugs) [1, 2]. They metabolically activate parent compounds to electrophilic intermediates, while Phase II enzymes conjugate these intermediates towards more easily excretable derivatives.

*CYP450* genes are a super-family of genes which appeared more than 3.5 billion years ago [3], being present in fungi, plants, bacteria, animals and humans. Genes are grouped into families and subfamilies based on sequence similarity: genes from the same family have sequence similarity greater than 40 % and, to be grouped into a subfamily, their sequence similarity must be greater than 55 % [4].

In humans, the *CYP450* family includes 57 genes and 58 pseudogenes [5] grouped in 18 families [6]. Several *CYP450* genes are found in clusters in the human genome but some members of the subfamilies can be spread out across the genome. For example, the CYP4F subfamily has genes on chromosome 19 and pseudogenes on multiple chromosomes. The *CYP2D6* gene is the most widely studied *CYP450* gene in humans, due to its role in the metabolism of many drugs [7, 8] along with *CYP3A4* and *CYP3A5*, members of the *CYP3A* subfamily[9, 10, 11, 12, 13]. However, not all *CYP450* genes or families have been studied thoroughly, and details on the evolution and clinical significance are lacking for several families, such as the *CYP4F* subfamily.

Several *CYP450* genes are potential candidates that underwent natural selection in humans [14, 15]. Other studies of the genetic diversity for specific *CYP450* subfamilies in human populations confirmed the presence of positive [16, 17], balancing [18] or purifying selection signatures [19]. One example is *CYP2C19*, involved in the metabolism of clopidogrel [20, 21], where signals of positive selection on its alleles conferring slow metabolism (*CYP2C19*\*2 and *CYP2C19*\*3) were detected using relative extended haplotype homozygosity (REHH)[18]. *CYP2C19*\*2 is detected worldwide, but *CYP2C19*\*3 is only present in people of Asian descent. The selective advantages may have been caused by diet and environmental pollutants impacting humans over thousands of years and could differ between ethnic groups. Additionally, low *F_ST_* values across *CYP2C19* SNPs suggest balancing selection in *CYP2C19* [18]. The excess of alleles at intermediate frequencies could reflect the evolution of balanced polymorphisms, which is to be expected in evolutionarily old enzymes responsible for numerous critical life functions.

Moreover, the detection of natural selection signals in the *CYP450* genes suggests that the selective advantage occurs at the molecular level. This selective advantage can act on polymorphisms that modulate gene expression, widely known as expression quantitative trait loci (eQTL) [22]. Detecting eQTLs linked to selection signals helps clarifying how gene expression is regulated and can lead to a better understanding of variants’ biological effects [23]. Furthermore, analysing eQTLs helps in the detection of gene-gene interaction [24] and co-regulation between genes [25]. Such genegene interactions can also be detected by looking at patterns of linkage disequilibrium (LD), as evolution will maintain co-evolving polymorphisms on the same haplotypes [26], which can also be detected as balancing selection signatures.

Here, we investigated genetic diversity and selective pressures across human populations in *CYP450* genes. Two subfamilies stood out in our analyses and were investigated in greater depth: the *CYP3A* and *CYP4F* families. We found that both families exhibit selective pressures in human populations and that the SNPs under selection could impact gene expression levels in several tissues. Furthermore, our results suggest interactions between the genes in both *CYP450* subfamilies, providing evidence of co-evolution and co-regulation within these gene clusters, that may vary between populations.

## 3 Results

We obtained genotypic data from the 1000 Genomes project phase 3 release[27] (1000G). A total of 2,157 individuals were analyzed from 22 populations, which were grouped into 4 super-populations (ie. Africa, Europe, East Asia and South Asia).

### 3.1 Global genetic diversity across populations in CYP450 genes

First, we aimed to identify global genetic patterns by calculating Tajima’s D values for each *CYP450* genes in each population of the 1000G dataset to provide insights into the non-neutral forces that act on these genes. A total of 61,739 biallelic SNPs were analyzed in the *CYP450* genes, and for each gene, we computed the mean Tajima’s D per gene and also in 1 Kb windows. Significantly low Tajima’s D values indicate an excess of rare alleles, whereas significantly high values of Tajima’s D suggest an excess of intermediate frequency alleles, which can reflect the occurrence of balancing selection.

In European populations, nine genes had Tajima’s D values consistently below 0 (Figure 1A). We assessed significance based on the empirical (null) distribution, which allows to determine whether any genes have values that are higher or lower than expected while taking population-specific demographic factors into account (see Methods). The proportion of 1 Kb-windows of each gene lying outside the null distribution is shown in Figure 1B. *CYP26A1, CYP27B1* and *CYP1A2* had the largest proportion of windows with significantly low D values, however these genes are quite small (4.4, 4.9 and 7.8 Kb, respectively), meaning that the signal is driven by one or two windows only. Interestingly, the four CYP3A genes in our dataset were all included in this group of nine genes, suggesting that strong purifying selection pressures may be acting, however complete selective sweeps driven by positive selection can also create this lack of diversity [28]. Notably, CYP3A5 has a low Tajima’s D average but no 1 Kb-window is significantly lower than expected, whereas other CYP3A genes have several windows showing significantly low Tajima’s D values. All *CYP450* genes show negative Tajima’s D values, as expected in coding regions, but ten genes have a mean above 0, which suggests relaxation of purifying selection pressure. The presence of several 1 Kb-windows significantly enriched for high D values can also reflect the presence of localized balancing selection signatures within these genes. Of these ten genes, five are in the CYP4F subfamily: *CYP4F3, CYP4F11, CYP4F12, CYP4F8* and *CYP4F2*. The strongest of these signals is seen on *CYP4F12* (Figure 1B). Interestingly, the only CYP4F gene that does not show this specific signature is *CYP4F22*, which is the ancestral gene of the *CYP4F* cluster [29].

**Figure 1:**
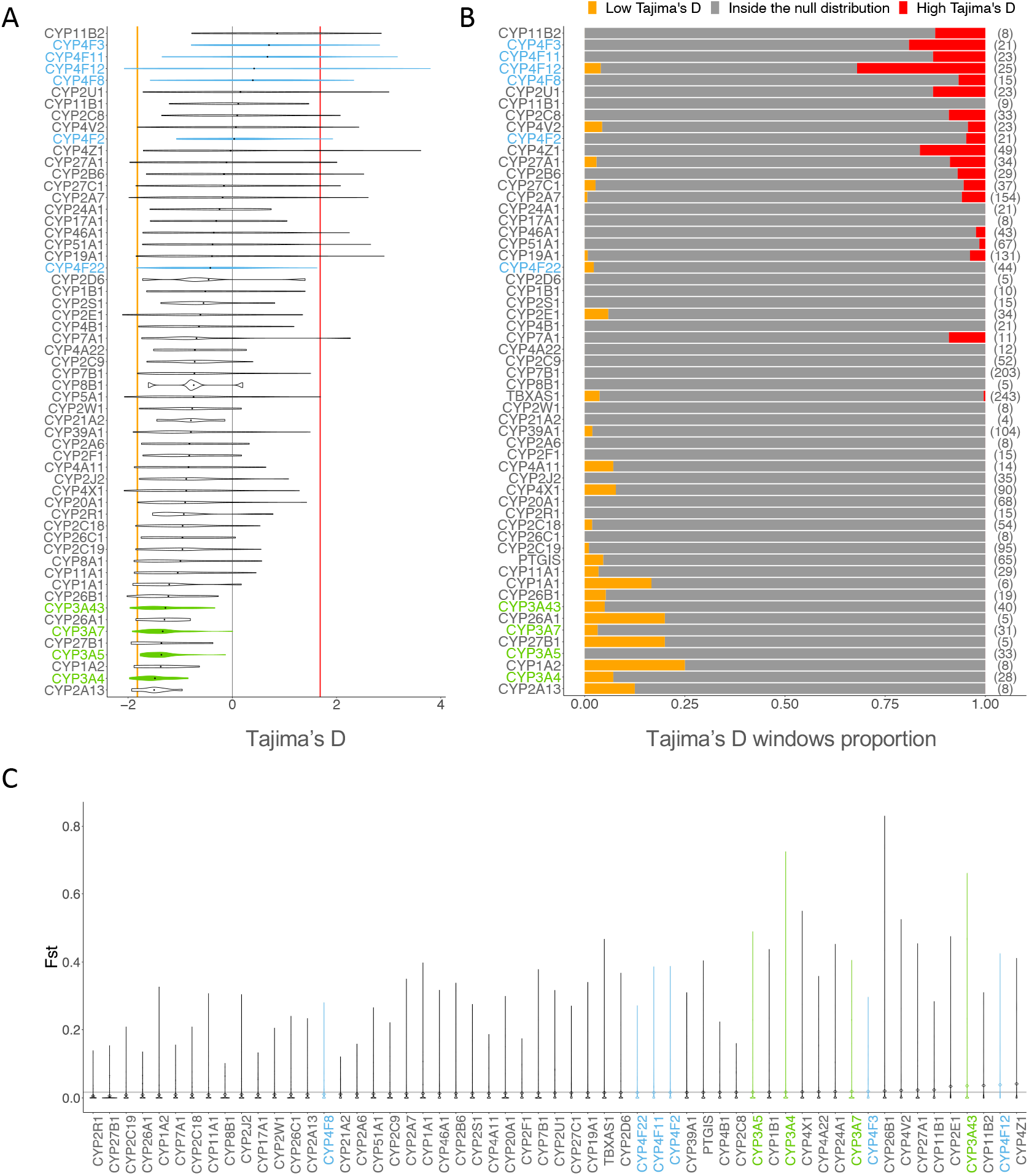
A) Distribution of Tajima’s D values computed on windows of 1 Kb for each CYP450 genes in the European populations. The 2.5th percentile is marked by the orange vertical line and the 97.5th percentile is marked by the red vertical line, representing the significance threshold. B) Proportion of Tajima’s D windows lying outside the null distribution for each CYP450 gene. For each gene, the total number of windows of Tajima’s D is shown beside the proportions, between brackets. The windows with Tajima’s D values below the 2.5th percentile is displayed in orange and over the 97.5th percentile is displayed in red. C) Distribution of *F_ST_* values for each CYP450 gene calculated on 4 super-populations (AFR, EUR, EAS, SAS). The mean *F_ST_* of chromosome 22, the null distribution, is displayed with the grey horizontal line.

Because population differentiation can also help identifying natural selection signatures within genes, we calculated the mean fixation index (*F_ST_*) across *CYP450* genes (Methods). *F_ST_* measures the differentiation between populations using genotype frequencies, with high *F_ST_* values indicating that the heterozygosity differs between populations. Figure 1C) shows the distribution of *F_ST_* values for each *CYP450* gene calculated on 4 super-populations (AFR, EUR, EAS, SAS). CYP4F genes are scattered across the *CYP450* spectrum, with *CYP4F12* having the second highest mean *F_ST_* while *CYP4F8* is in the bottom half of the distribution. Mean *F_ST_* of genes of the *CYP3A* subfamily are in the highest values, meaning that these genes have a high divergence between population’s genotype frequencies. This could indicate that the low Tajima’s D in *CYP3A* reflects positive rather than extreme purifying selection.

### 3.2 Positive selection in *CYP3A* and *CYP4F* subfamilies

The global neutrality and differentiation analyses of *CYP450* genes suggest that positive selection, either directional (*CYP3A*) or balancing (*CYP4F*), may be acting on subfamilies of *CYP450* genes, possibly in a concerted fashion. To further validate positive selection signatures and identify specific putative sites, we used the integrated haplotype score (iHS), which leverages linkage disequilibrium (LD) patterns in a specific population [14]. Typically, an absolute value of iHS greater than 2 at a SNP suggests that the region around the SNP is under selection [14].

In the *CYP3A* cluster, significant iHS values are detected (Figure 2A), but signals of positive selection differ between populations. Many signals are detectable in Africans, in East Asians and in Europeans, while fewer signals are detectable in South Asians. Signals of positive selection are noticeable in *CYP3A5, CYP3A51P, CYP3A4* and *CYP3A43* among Africans. In particular, iHS values in *CYP3A5* are consistently below −2, indicating that the derived alleles have quickly increased in frequency, a signature of positive selection. Among East Asians, the selective sweep is located from *CYP3A51P* to *CYP3A4*, and among South Asians, in *CYP3A43*. Lastly, for Europeans, signals of positive selection are detectable in the region between *CYP3A7* and *CYP3A4*, a signal also present in the East Asian population. *CYP3A43* is the only gene with signals in all super-populations. These results confirm that positive selective pressure is acting on *CYP3A* genes.

**Figure 2:**
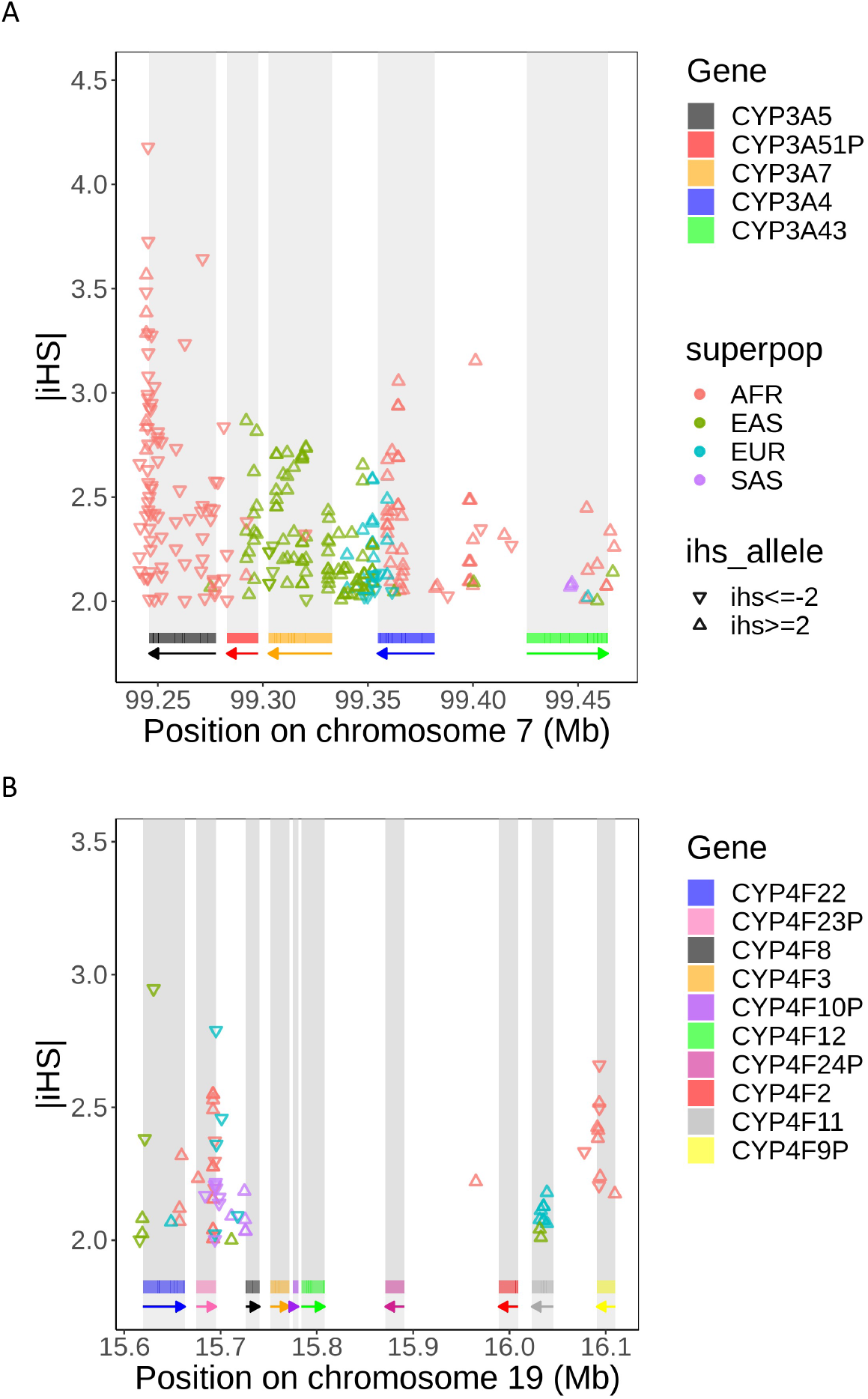
Distribution of SNPs with high |*iHS*| values (|*iHS*| ≥ 2) in the A) CYP3A and B) CYP4F cluster. A triangle standing on its base means an *iHS* value ≥ 2, indicating that the ancestral allele has increased in frequency, and a triangle standing on its point means an *iHS* value ≤ −2, indicating that the positive selection is acting on the derived allele. SNPs located in repetitive elements and sequences are masked. Rectangles below the plot show the position of each gene and arrows indicate on which strand the gene is located.

Positive selective pressure is also detected in the *CYP4F* cluster, but on a smaller scale. For the *CYP4F* cluster, signals of positive selection are visible in *CYP4F22*, *CYP4F23P*, *CYP4F11* and *CYP4F9P* (Figure 2B). The region between the pseudogene *CYP4F23P* and the gene *CYP4F8* also shows high iHS values, indicating positive selection in every super-population. iHS values greater than 2 are present in *CYP4F11* in Europeans and East Asians, indicating positive selection acting on ancestral alleles. *CYP4F9P* has significant iHS values in Africans. Again, most iHS values are greater than 2, indicating selective pressures on ancestral alleles, but the 3 strongest signals are seen for derived alleles (iHS below −2), suggesting these SNPs may be driving the signal.

### 3.3 Balancing selection in *CYP3A* and *CYP4F* subfamilies

The Tajima’s D analyses (Figure 1) suggested balancing selection in the *CYP4F* cluster. To confirm this finding, we used the Beta score [30], a statistic which detects clusters of alleles with similar allele frequencies, developed to specifically test whether balancing selection is present at specific loci.

We considered *β* score in the top 1% of the whole chromosome as significant *β* scores (empirical p-value < 0.01), which can vary between populations. In contrast to iHS, very few significant *β* score values are seen in the *CYP3A* cluster. Only one SNP in *CYP3A43* meets this criteria in Africans. The same signal can been seen in the other populations, but it is weaker and do not pass our 1% threshold (Figure 3A). Overall, these results show no clear evidence of balancing selection acting on the *CYP3A* cluster. In line with Tajima’s D results, clearer signals are seen in the *CYP4F* cluster, which show larger *β* scores compared to the *CYP3A* cluster: the highest *β* score in the *CYP4F* cluster is almost twice as high as the highest *CYP3A*’s *β* score. SNPs in *CYP4F12* show highly significant *β* scores, replicated among Africans, Europeans and South Asians, but not in the East Asians. Also, the region between *CYP4F23P* and *CYP4F8* has the most extreme *β* score in the region, and the signal is visible in all super-populations (Figure 3B). These consistent signals across populations provide convincing evidence of balancing selection acting around *CYP4F8* and *CYP4F12*. Weaker signals, which do not pass our significance threshold but are seen consistently between populations, are seen in *CYP4F23P* and *CYP4F11*. Taken together, these results confirm that balancing selection does act on the *CYP4F* cluster, but not in the *CYP3A* cluster.

**Figure 3:**
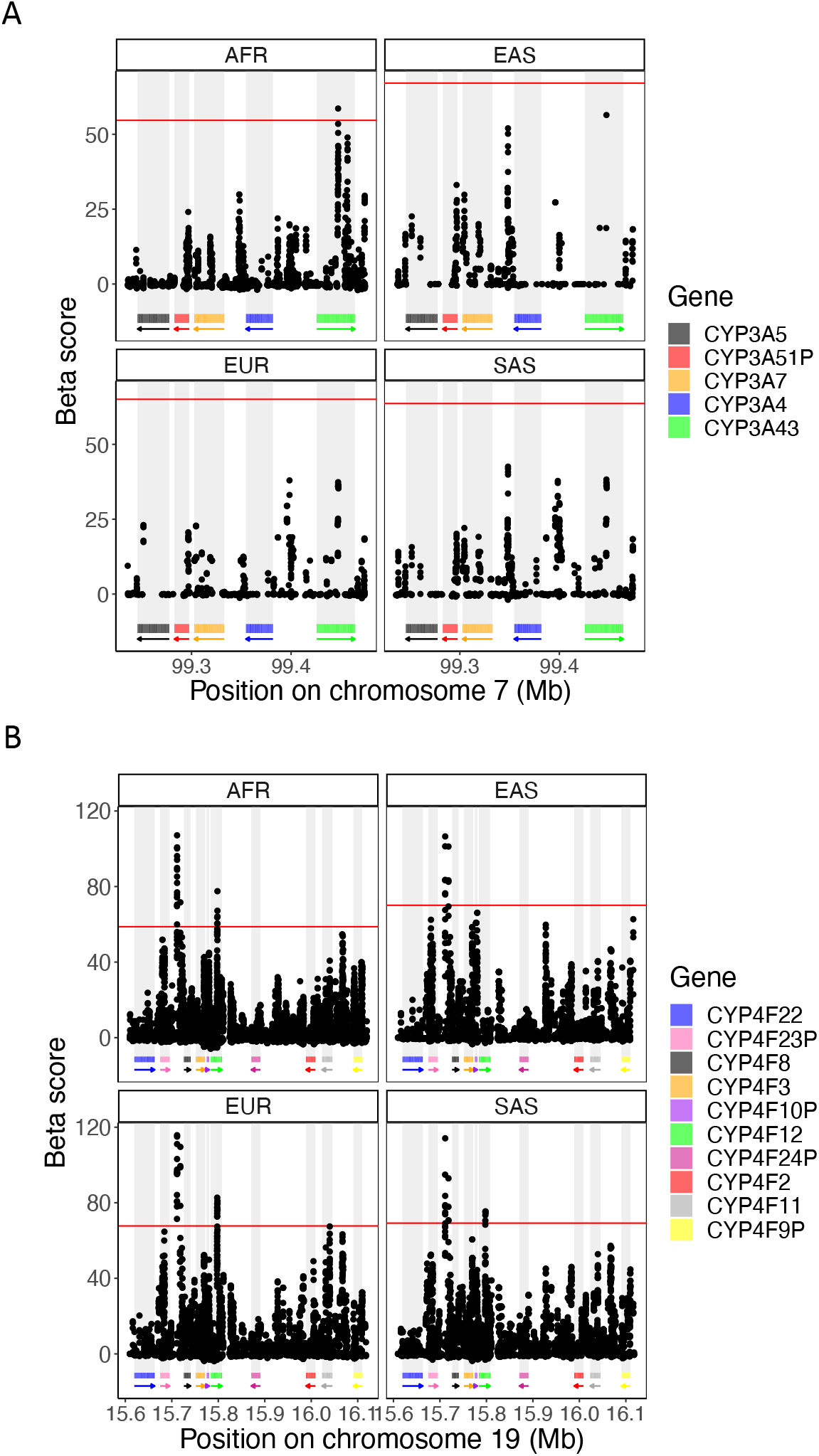
*β* score in the chromosomal region of the A) CYP3A and B) CYP4F cluster for the 4 super-populations analyzed. The *β* score was calculated on the 1000G dataset and the 99th percentile indicating the top 1% *β* score is displayed by the horizontal line in red. Rectangles below the plot show the position of each gene and arrows indicate on which strand the gene is located.

### 3.4 Detection of Unusual Linkage Disequilibrium

Since *CYP3A* and *CYP4F* genes are in a gene cluster and selective pressures are acting on these genes, co-evolution could be occurring. Indeed, the different combinations of alleles which co-occured during evolution can lead to concerted selective pressure, or co-evolution, depending on the resulting fitness of the individuals [26]. Such co-evolution signals can be revealed by analyzing patterns of linkage disequilibrium (LD) beyond local associations due to allelic proximity, in order to detect whether specific combinations of alleles (or genotypes) at two distinct loci are particularly overrepresented. To do so, we calculated the genotyped-based LD (r^2^) between each pair of SNPs with minor allele frequency (MAF) above 0.05 in the two CYP450 clusters, across each 1000G subpopulation (Methods). Under neutrality, the LD association between SNPs is expected to decrease as genetic distance between the SNPs increases, allowing us to build an empirical distribution by considering clusters of genes of similar size genome-wide (Methods) to the clusters under investigation. Pairs of SNPs showing unusual LD (uLD) values, lying outside of this null distribution, are therefore likely transmitted together more often than expected, making it possible to identify candidate sites that are co-evolving.

In both clusters, strong signals of uLD are present (Figure 4, Figure S1) compared to matched gene clusters (Methods), with *CYP4F* showing much more extreme signals than *CYP3A* (8.1% vs 4.7% of pairs of SNPs in uLD), despite genetic distances in the *CYP4F* cluster being four times larger than in the *CYP3A* cluster (maximum distance of 0.60 cM vs 0.15 cM, respectively), whereas the physical size of the cluster is only twice (500 Kb vs 250 Kb, respectively). Significant uLD between *CYP3A5* and *CYP3A43* and between *CYP3A7* and *CYP3A43* can be seen in all European populations (Figure S1A). *CYP3A5* and *CYP3A43* are the opposite to each other in term of physical location in the cluster while *CYP3A7* and *CYP3A43* are next to each other. Finland (FIN) and Toscani (TSI) populations have the most uLD signals across European populations, with FIN uniquely showing uLD between *CYP3A5-CYP3A4*, and TSI showing uLD between *CYP3A4* and *CYP3A43*, a signal consistently seen in the East Asians. TSI also have the highest genetic distance interval in this region, likely due to a larger, more widespread, recombination rate in *CYP3A4* compared to other populations (Figure S2). Among East Asians, uLD signals are seen almost exclusively between SNPs in *CYP3A4* and *CYP3A43*, two genes that are next to each other, with no clear recombination hotspot separating them, meaning that linkage disequilibrium can be expected (Figure S2). SNPs in these genes are also in uLD in Gujarati Indian (GIH) population, but none of the other South Asian populations show any signal, which may be explained by the short genetic distances within this cluster in this super-population (SAS) (<0.05 cM). Finally, African populations show the most deviation from the null (Figure 4A). SNPs in *CYP3A4* are in uLD with all other genes and the signal also replicates the observations from the European populations, with SNPs in *CYP3A43* in uLD with SNPs in all other genes (Figure 4A, Figure S1A).

**Figure 4:**
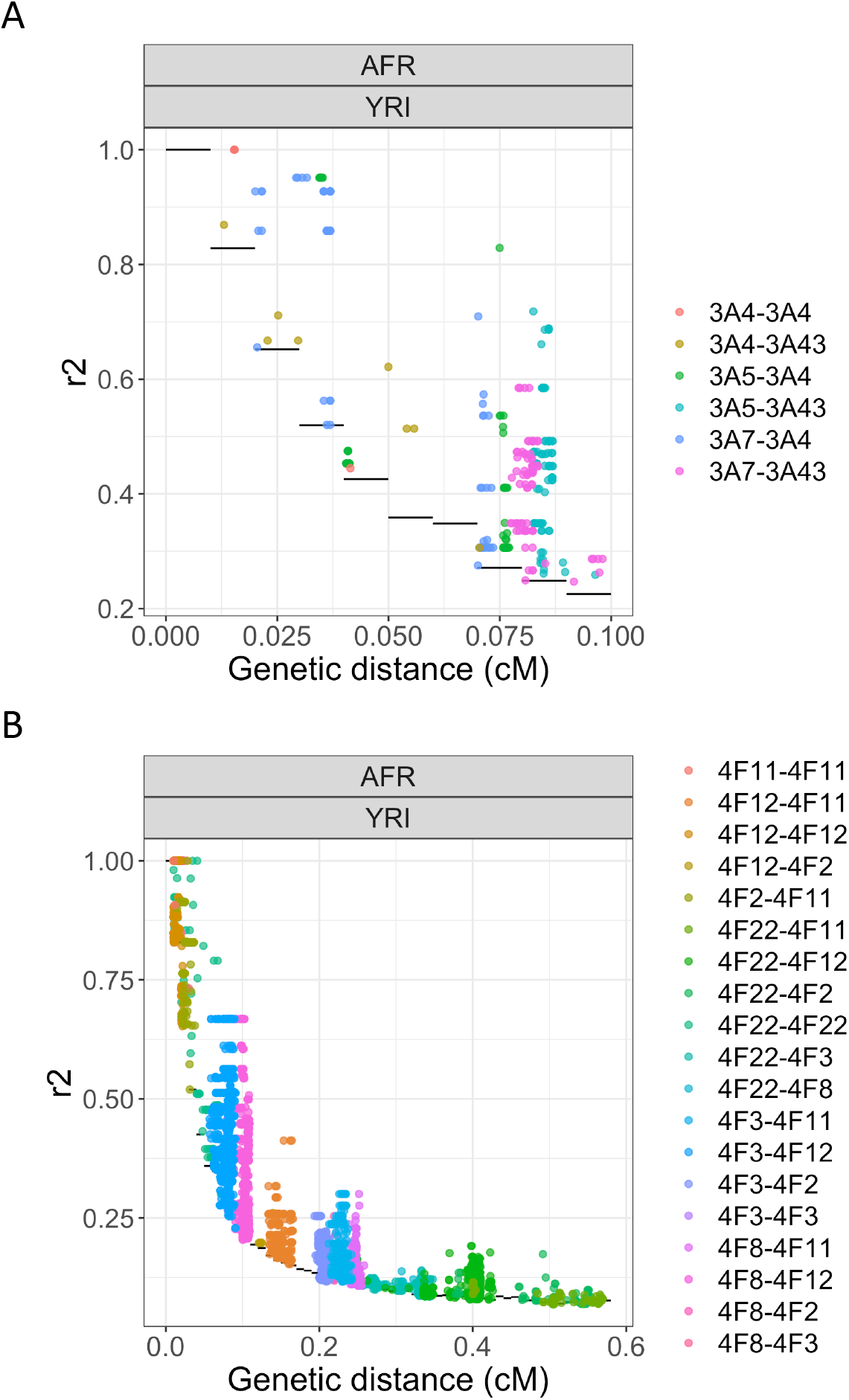
r^2^ values between each pairs of SNPs in the A) CYP3A and B) CYP4F cluster in the Yoruba (YRI, AFR) population. The distance between the SNPs is in centimorgan (cM). Only r^2^ values over the null distribution are shown. The null distribution is shown with black horizontal lines. Dots are colored according to which genes are involved in the pair.

In the *CYP4F* cluster, several pairs of SNPs have patterns of LD that deviate significantly from the empirical distribution (Figure S1B). There is uLD for *CYP4F22-CYP4F11* and *CYP4F22*-*CYP4F12* in almost every populations, even though these genes are far from each other (0.36 Mb and 0.12 Mb, respectively). *CYP4F22* and *CYP4F2* are also in uLD in AFR, EUR and EAS.

The African populations have more evidence of uLD than the other super-populations. One population in particular, the Yoruba (YRI) population, has even more extreme signals in comparison with other African populations and most uLD signal are driven by associations involving the *CYP4F12* gene (Figure 4B). Thus, we investigated whether a specific region in *CYP4F12* is in strong LD with the other genes. Indeed, in the YRI population, there is evidence of uLD between a region in *CYP4F12* (at 15.79 - 18.00 Mb on chromosome 19) and the *CYP4F3* (Figure S3A) and *CYP4F8* genes (Figure S3B). The extreme signals in this gene cluster are in line with the hypothesis that balancing selection acts via gene-gene interactions, or epistasis [31]. As these patterns could be due to sequencing errors [32], we used the latest 1000G dataset which has high-coverage sequencing and is aligned on GRCh38 (Methods). These results were replicated in this second dataset, greatly reducing the possibility that the observed signal is due to sequencing errors or spurious mapping. Finally, in the Europeans, the FIN population has a specific pattern between *CYP4F12* and *CYP4F2*, *CYP4F8*, *CYP4F3*. Looking more closely, many SNPs in *CYP4F12* are in uLD with one SNP in *CYP4F3* (Figure S3A) and two SNPs in *CYP4F8* (Figure S3B). No specific SNPs are in uLD with *CYP4F2*.

### 3.5 Detection of eQTLs

We next evaluated the effects of the SNPs identified as being under positive and balancing selection on the expression of the genes in each *CYP450* cluster to test if these are eQTLs.

In the *CYP3A* cluster, three SNPs are under positive selection in the Punjabi population from South Asia (PJL): rs487813, rs679320 and rs568859. These SNPs are located in *CYP3A43* and are significant eQTLs of *CYP3A5* in lung (Figure 5A). The SNP under balancing selection in the Luhya population (LWK) in *CYP3A43*, rs800667, is also an eQTL of *CYP3A5* in lung (Figure 5B). The effect size estimate for these significant eQTL is negative, indicating a reduction in *CYP3A5* gene expression with each non-reference allele. This locus in *CYP3A43* thus impact *CYP3A5* expression in lung, even though *CYP3A5* and *CYP3A43* are at opposite ends of the cluster, 147.99 Kb apart. This result is in line with the LD analyses (Figure 4A), which suggested uLD between SNPs in *CYP3A5* and *CYP3A43* in Europeans, Africans and the Japanese. Those four SNPs were all in uLD with 11 SNPs in the Toscani population (TSI) and five other SNPs in Americans of African Ancestry (ASW).

**Figure 5:**
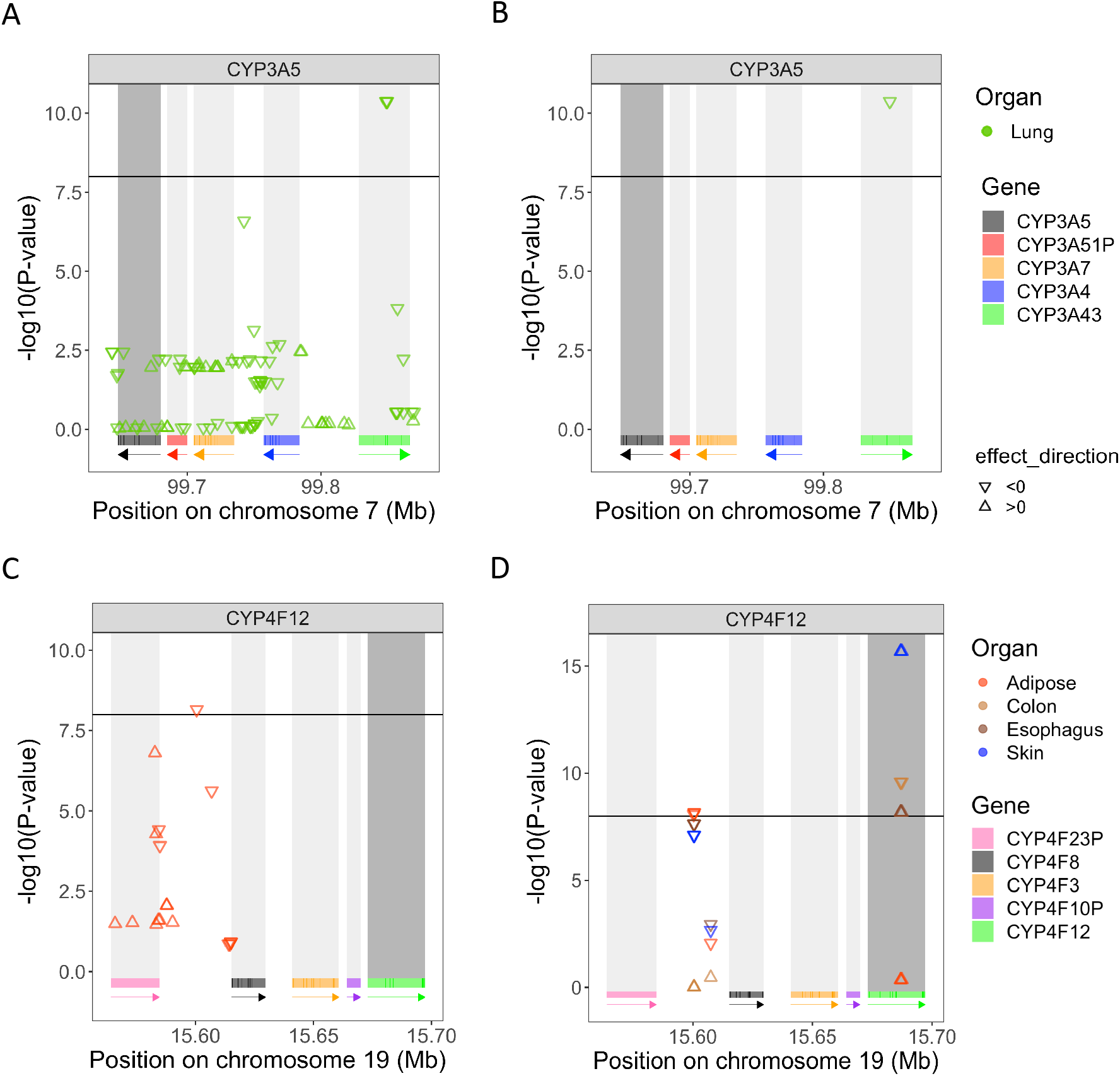
P-values of the associations between SNPs under A) positive selection and B) balancing selection and CYP3A5’s gene expression in lung and p-values associated with SNPs C) under positive selection and D) balancing selection and tissue-specific gene expression of CYP4F12. CYP3A5 and CYP4F12 are shown in dark gray, as the expressions of these genes are tested. The triangle standing on its base indicates a positive effect size (*β_eQTL_* > 0), while a triangle standing on its point indicates a negative effect size (*β_eQTL_* < 0). The threshold, set to 10^-8^, is represented by the horizontal black line, meaning that a −*log10*(*p* – *value*)>8 is a significant eQTL. Only tissues with significant eQTLS are displayed. As before, rectangles below each plot show the position of each gene and arrows indicate on which strand the gene is located. Each gene has its own colour to indicate its location.

In the *CYP4F* cluster, a SNP under positive selection, rs74459786 (Table S1), located in the intergenic region between *CYP4F23P* and *CYP4F8*, is an eQTL of *CYP4F12* in adipose tissue (Figure 5C), with a negative effect size. SNPs under balancing selection (Table S2) within *CYP4F12* are eQTLs for *CYP4F12* expression in the colon, esophagus and skin, but interestingly, their effects in these tissues are in opposite directions, with positive effect sizes in the colon and skin, and negative ones for the esophagus. Furthermore, a SNP with a balancing selection signal is also an eQTL of *CYP4F12* expression in adipose-subcutaneous tissue (Figure 5D) with a negative effect size estimate. It lies in the intergenic region between *CYP4F23P* and *CYP4F8*, which is the same region as the SNP under positive selection (rs74459786) in Figure 5C.

Another SNP under positive selection in this intergenic region, rs62115147 (Table S1), is also associated with *CYP4F3* expression in one of the brain tissues (Brain-Spinalcord-cervicalc-1) and in nerve tissue (Figure S4A). The *CYP4F12* gene emerged repeatedly as a candidate in our balancing selection and uLD analyses, while the intergenic region between *CYP4F23P* and *CYP4F8* is seen only in the balancing selection analysis.

Even if less positive selection is present in the *CYP4F* cluster compared to the *CYP3A* cluster, many of the SNPs showing high iHS values in the *CYP4F* cluster show up as eQTLs for different genes. SNPs under positive selection located in *CYP4F11* (Table S1) are eQTLs of *CYP4F2* in brain and skin tissues (Figure S4B) with consistent, negative effect sizes. Additionally, the same SNPs under positive selection within *CYP4F11* are associated with expression of *CYP4F11* itself in multiple tissues (Figure S4C). The direction of effect on gene expression is the same for all significant associations.

### 3.6 Phenotypic associations

Using the UK Biobank cohort (UKb), we did a Phenome-Wide Association Study (PheWAS) to identify phenotypes potentially under selective pressure (Methods), using the available variants with selective signals in the *CYP4F* genes (166 variants from the 180 found under selection) and in the *CYP3A* genes (62 from the 125 variants found under selection). No significant associations were found for SNPs under selective pressure in *CYP4F* cluster.

In the CYP3A cluster, however, SNPs under positive selection in at least one studied populations were found associated with six phenotypes (Figure 6A,B) in our PheWAS. Among the disease phenotypes, we found association with pelvic inflammatory disease (PID), which is female-specific, and for which the SNP with the strongest association (p-value_*rs*2014764_ = 1.96×10^-5^) was under positive selection in European (CEU, GBR) and East Asian (CHB, CHS, CDX) populations. Among the continuous phenotypes investigated (Methods), we found association with pulse rate, for which the SNP with the strongest association (p-value_*rs*12536946_ = 4.66×10^-13^) was also found under selective pressure in Europeans (CEU). Among the biomarker variables, the strongest associations with platelet count (p-value_*rs*503115_ = 2.30×10^-12^) and erythrocyte count (p-value_*rs*10235630_ = 1.83×10^-7^) were both found with SNPs under selective pressure in the Japanese population.

**Figure 6:**
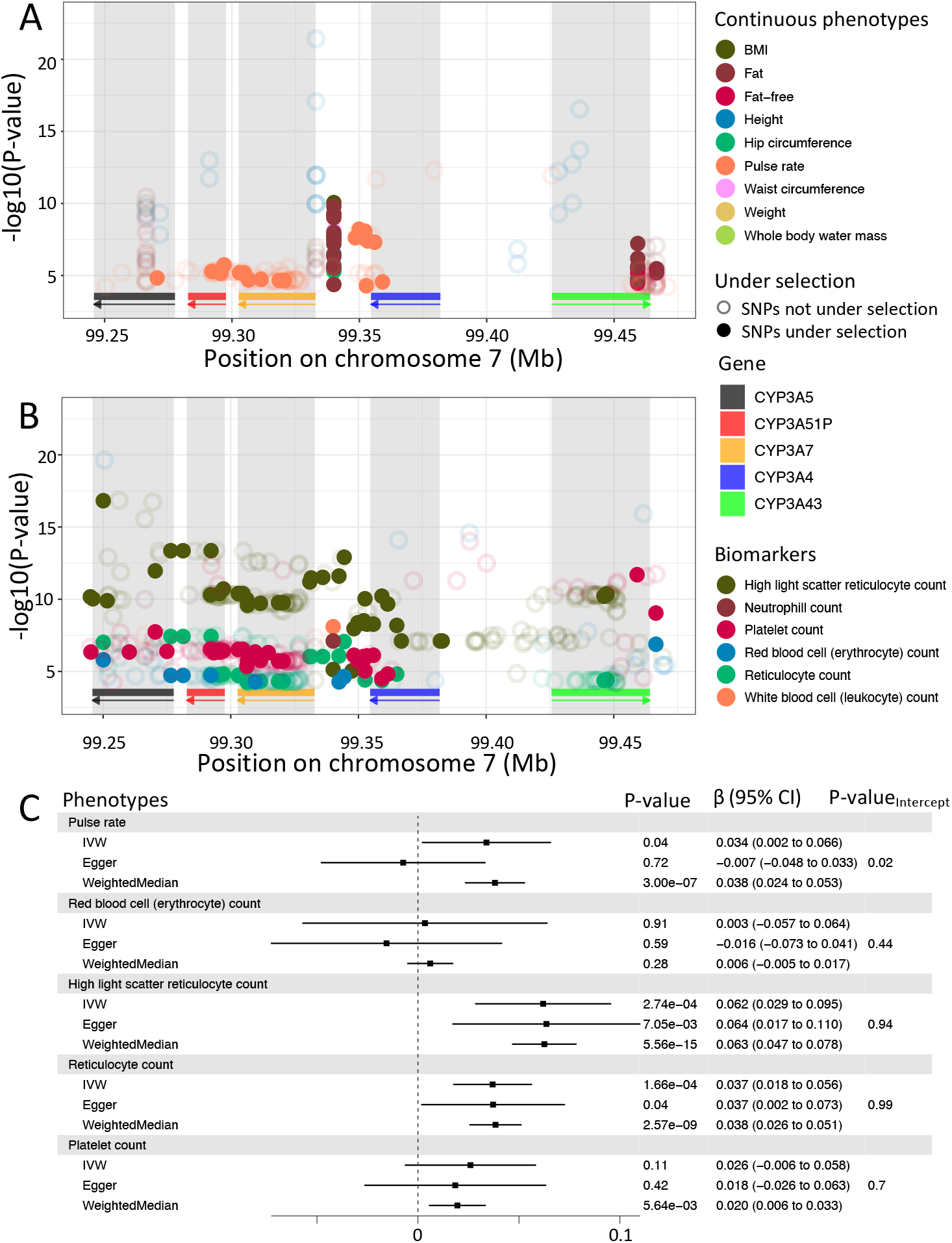
Associations of CYP3A cluster with phenotypes in the UK biobank. Significant associations (p¡0.05/777) for continuous traits (A) and plasmatic biomarkers (B) which are significant in at least one SNP under selection. SNPs under selection are represented as full dots, meanwhile other SNPs are represented as empty dots. As before, rectangles below each plot show the position of each gene and arrows indicate on which strand the gene is located. Each gene has its own colour to indicate its location. C) Causal relationship with CYP3A5 expression in lung for phenotypes showing significant association with its eQTLs. *β* represents the change of 1 standard deviation of CYP3A5 expressions on phenotypes, also in standard deviation units. P-value of three statistics (IVW, Egger, Weighted Median) are displayed with the *β* and the 95% confidence interval (CI) of the association for each phenotype in the grey box. For Egger, the p-value of the intercept is also displayed.

Lastly, for both high light scatter reticulocyte count and reticulocyte count, their strongest association (p-value_*rs*73713580_ = 1.24×10^-17^; p-value_*rs*55830753_ = 3.08×10 ^8^ respectively) were both found under selective pressure in African population (MSL and ACB, respectively). Using Mendelian randomisation (Methods), we evaluated the causal relationship between *CYP3A5* expression in lung, for which eQTLs were found under selective pressure above, and the phenotypes found to be associated with SNPs in the *CYP3A* cluster. We identified a significant causal association between CYP3A5 expression and both high light scatter reticulocyte count (p-value_*IVW*_ = 2.74×10^-4^) and reticulocyte count (p-value_*IVW*_ = 1.66×10^-4^). We did not detect pleiotropy using MR-Egger and results were robust using the weighted median test (Methods).

Altogether, these results indicate that the selective pressure in the *CYP3A* cluster could be driven by the production of reticulocyte through the expression levels of *CYP3A5*, and also suggest that pulse rate could be impacted by genetic variation in *CYP3A* genes.

Among other associations identified in this cluster, three SNPs showed strong associations with anthropometric traits (Figure 6A) and are under selective pressure in European population (CEU, IBS). Those SNPs were, however, found to be associated with expression of genes outside the *CYP3A* cluster (Supplementary text 10.2), prompting for further investigation of the relationship between this cluster and other neighbouring genes to understand the different drivers at play.

## 4 Discussion

Drug metabolism is a rather complex system modulated by many *CYP450* genes. As shown by others [17, 33, 34, 35], we also found that selective pressure and genetic differentiation between populations were present in *CYP450* genes. Here, we provide a deeper analysis of two *CYP450* clusters, the widely studied CYP3A [34, 35, 36, 37] and the less well-known *CYP4F* clusters, identified thanks to their outlier patterns in neutrality and population differentiation analyses. These two *CYP450* clusters have higher levels of natural selection forces (positive selection and balancing selection) acting on them as a whole and show population differentiation. We found that natural selection forces involved differ between the two clusters; the *CYP3A* cluster is evolving under positive selection, while the *CYP4F* cluster show signals of balancing selection. Furthermore, the *CYP4F* cluster shows strong evidence for co-evolution and co-regulation signals.

In the literature, the *CYP450* genes are often studied independently. In our study, genes are studied as families, and we determined that there is evidence of epistasis between the genes within each cluster. As these clusters of genes are involved in drug metabolism [1, 2, 38, 39], it is important to understand the impact of genetic variants on their gene expression, to help understand how these variants might impact drug response and refine disease treatments in a personalized way. Our results also show that the impact of specific variants may differ between populations, which could lead to a deeper understanding of differences in individual drug response [40, 41].

The *CYP3A* cluster contains 4 genes: *CYP3A4, CYP3A5, CYP3A7* and *CYP3A43*. Signals of positive selection were detected in the *CYP3A* cluster, specifically in *CYP3A4* and *CYP3A7*, which have been under recent positive selection in African, European and the Chinese populations, while *CYP3A5* appears under positive selection in Europeans and *CYP3A43* in non-Africans [35]. Our analyses confirmed that *CYP3A* genes are evolving under positive selection as previously reported [14, 17, 33, 34].

We found that the locus known to cause non-expression of *CYP3A5* [42], rs10264272/ *CYP3A5*\*6, is under positive selection | ≥ 2) in African populations (YRI, GWD, LWK). A second locus, known to cause low *CYP3A5* expression, rs776746/ *CYP3A5*\*3, is under positive selection (|*iHS*| ≥ 2) in two African populations (YRI, GWD). These derived allele have thus swept up in frequency in several African populations. In the Toscani population, rs776746/*CYP3A5*\*3 is found to be in uLD with the four SNPs under selective pressure in the *CYP3A* cluster, that are eQTLs of *CYP3A5* in lung.

*CYP3A43* is the ancestor gene of this cluster [17, 43], however, its function is not well understood, unlike other *CYP3A* genes. Our analyses suggest that SNPs in *CYP3A43* regulate *CYP3A5* gene expression, at least in lung. Levels of expression of *CYP3A5* in lung were causally associated to reticylocytes count and many of its eQTLs were under selection in Africans. Since *plasmodium vivax*, a parasite causing malaria, affect mainly young reticulocytes [44] and that malaria is present in Africa, the selective pressure found in this population could be associated to this disease. Further studies need to be done to validate this hypothesis.

In the *CYP4F* cluster, we found both positive and balancing selection pressures acting. Furthermore, the SNPs evolving under selective pressures are associated with gene expression levels across the cluster in several tissues. For instance, a cis-eQTL of *CYP4F12*, rs74459786, is detected to be under positive selection in the Kinh population in East Asia (KHV). We also found that several SNPs in *CYP4F11* are associated with *CYP4F2* expression. Both genes are implicated in common metabolic function, such as the synthesis of 20-HETE (20-hydroxyeicosatetraenoic acid) from arachidonic acid [45]. Thus, this could indicate a possible regulatory mechanism of common functions.

Finally, the region between *CYP4F23P* and *CYP4F8* emerged multiple times in our analyses. This intergenic region shows strong signals for selection, with the same SNPs also being eQTLs of *CYP4F3* (nerve) and *CYP4F12* (adipose tissue). Given the implication of *CYP4F12* in fatty acid metabolism [46], our results may point towards the identification of new regulatory elements involved in this process in adipose tissues.

In conclusion, our results demonstrate high heterogeneity across human populations, both in terms of selective signals and interaction between variants and expression levels, for the *CYP3A* and *CYP4F* genes. There could thus be important differences in metabolic regulation impacting drug response in individuals from different ethnicities. In particular, these variants could cause impaired efficacy, as well as side effects. As pharmacogenetic studies still typically focus on European populations, our results underline the importance of including individuals from several populations in order to capture all of the genetic diversity and its impact on disease treatment and metabolism.

## 5 Methods

### 5.1 1000 Genomes genetic data

The data analyzed is from the Phase III of the 1000 Genomes project (1000G) [27]. The 1000 Genomes Project includes 2,504 individuals from 26 populations. These populations can be split into 5 super-populations: African (AFR), European (EUR), South Asian (SAS), East Asian (EAS) and Admixed American (AMR). Data from the AMR population is not included in this study because the high degree of admixture may confound selection and linkage disequilibrium analyses. This left us with 22 sub-populations and four super-populations for study. The available variant call format (vcf) files of 1000G are under the GRCh37 genome build. VCFtools v0.1.14 [47] was used to filter the 1000G dataset. Indels and non-biallelic alleles were removed and only SNPs located in the 57 CYP450 genes were kept, extracted based on coordinates from the UCSC Genome Browser.After filtering, the CYP450 dataset included a total of 61,739 SNPs and 2157 individuals. We refer to this as the “1000G CYP450 dataset”. A more recent dataset was also used, the re-sequencing dataset of 30X coverage, mappend on GRCh38 [48], which includes the 2157 individuals.

### 5.2 Genetic diversity and population differentiation

Both Tajima’s D and *F_ST_* statistics were obtained with VCFtools using the 1000G CYP450 dataset. Tajima’s D values were calculated in the super-population (AFR, EUR, EAS and SAS) separately on non-overlapping windows of 1 Kb. We computed the mean Tajima’s D value for each gene by averaging the window-based values, and sorted genes according to their mean. To create a null distribution, we computed Tajima’s D values for all SNPs associated with a gene name in the CADD annotation file on chromosome 22, so that all SNPs used to compute the empirical distribution are located in genes. We computed the 2.5 and 97.5th percentile on the window-based values of chromosome 22. Values above the 97.5th percentile and below the 2.5th percentile were considered to be statistically significant (two sided empirical p-value <0.05). The *F_ST_* values, from Weir and Cockerham derivation [49], were calculated using four super-populations (AFR, EUR, EAS and SAS) on a per-site basis. The per-gene mean was calculated on raw values and genes were sorted based on their mean *F_ST_*. As in the previous analysis, chromosome 22 was used to create an empirical distribution. *F_ST_* values were also computed on SNPs located in genes of the chromosome 22 (see above) and the per-gene mean *F_ST_* was calculated.

### 5.3 Detecting natural selection

The method used to detect balancing selection is the *β* score [30]. This score has already been calculated on the whole 1000 Genomes project data. The approach used to detect signal of recent positive selection was iHS (integrated haplotype score) [14]. The iHS computation was performed by us on the 1000G dataset, filtered to exclude INDELs and CNVs. Reference alleles from filtered 1000 Genomes vcf files were changed to the ancestral alleles retrieved from 6 primates EPO pipeline (version e59) using the `fixref` plugin of bcftools [50]. The hapbin program v.1.3.0 [51] was then used to compute iHS using per population-specific genetic maps computed by Adam Auton on the 1000 Genomes OMNI dataset. When the genetic map was not available for a sub-population, the genetic map from the closest sub-population was selected according to their global *F_ST_* value computed on the 1000G dataset. For all natural selection analyses, SNPs annotated to be in a repetitive region were identified using the RepeatMasker track available on the UCSC genome browser [52] and were removed.

### 5.4 Unusual Linkage disequilibrium

Linkage disequilibrium between pairs of SNPs from the same cluster was assessed using the `geno-r2` option from VCFTools on SNPs with minor allele frequencies (MAF) above 0.05. The genetic position of each SNP was calculated with PLINK v1.90 [53] using the population-specific genetic maps the same was as described in previous section.

To compute a null distribution to detect unusual linkage disequilibrium (uLD), the Human GRCh38 Gene transfer format (GTF) file from Ensembl v87 was screened per chromosome using an in-house python script to find windows matching the CYP4F cluster: windows of 430 Kb containing 6 genes were kept. In these windows, we excluded INDELs and SNPs with MAF <0.05. The r^2^ for each pair of SNPs located within a selected window was computed using VCFTools with the `geno-r2` option. We divided the genetic distance into bins of 0.01 cM and wcalculated the 99th percentile of *r*^2^ values of each pair of SNPs lying in the bin. This process was done separately for each 1000G sub-population, yielding a null distribution per sub-population. r^2^ values on pairs of SNPs in the extremes of the empirical distribution are considered to be significant for what we called unusual linkage disequilibrium (uLD).

To specifically confirm the signal seen between CYP4F12 and other CYP4F genes, we extracted only the SNPs showing significant uLD in the previous analysis and kept only those pairs where one SNP was located in CYP4F12. Using VCFTools, CYP4F genetic data was extracted from the 1000 Genomes 30X on GRCh38 dataset [48] and *r*^2^ values were calculated as described above.

### 5.5 eQTLs analysis of SNPs under selection

The Genotype-Tissue Expression v8 (GTEx)[54] was accessed through dbGaP (phs000424.v8.p2, dbgap project #19088) and contains gene expression across 54 tissues and 948 donors as well as genotyping information, compiled in a VCF file by GTEx on the GRCh38 genome build. The cohort comprises 67% males and 33% females, mainly of European descent (84.6%), aged between 20 and 79 years old. Analyses were done on 699 individuals of European descent, as described in Supplementary text (*Pre-processing of GTEx genetic data*). To take into account hidden factors, we calculated PEER factors on the normalized expressions. We removed tissues with less than 50 samples, leaving samples from 50 different tissues.

For eQTL analyses, we selected only SNPs that were identified to be under positive or balancing selection in CYP3A and CYP4F clusters in previous analyses. Since the positions of these SNPs were in the GRCh38 genome build, we converted these positions to the GRCh38 genome build to match GTEx v8 data, using the `liftOver` function of the `rtracklayer` R library [55]. P-values of associations between each selected SNP and gene expression of every gene in the cluster were calculated with a linear regression using the lm function in R. The linear regression was calculated on each SNP individually. The covariates include the first 5 principal components (PCs) (see Supplementary text), age, sex, PEER factors, the collection site (SMCENTER), the sequencing platform (SMGEBTCHT) and total ischemic time (TRISCHD). To report genome-wide significant eQTL signals, we used a p-value threshold for significance at 10^-8^.

### 5.6 Phenotypic associations

The UK biobank (UKb) [56] was accessed through project 15357. We kept only individuals of European descent which were within 3 standard deviation of the mean for the top 3 PCs, removed one individual for each pair of related individuals, and removed individuals whose genetic sex did not match self-identified sex. We extracted positions for CYP3A (chr7:99-244-812-99-470-881, GRCh38) and for CYP4F (chr19:15-618-335-16-110-830, GRCh38) families. We then removed positions with more than 10% of missing genotype and with a MAF under 1%, then removed individuals with more than 5% missing genotypes, leaving us with 399,149 individuals and 3,092 variants for the CYP4F genes, and 400,504 individuals and 374 variants for the CYP3A genes

We used baseline values for continuous phenotypes. We selected phenotypes recommanded by the UKb, as well as blood cells measurements, a total of 90 and 11 phenotypes respectively (Table S3). When many values were available at the baseline, we took the mean of those values. We also looked at diseases using phecode coding extracted from phewascatalog [57], which indicated ICD-10 to group and to exclude from controls. We kept only phecodes with more than 500 cases, leaving 603 for both sexes, 62 female-only and 11 male-only. Covariates used are the age at baseline, sex, top 10 PCs, deprivation index and the genotyping array. Analyses were done with plink2 [58] with linear transformation of the quantitative covariates. We used a p-value threshold for significance at 6.44×10^-5^, based on Bonferonni correction for the number of phenotypes evaluated (0.05/(90+11+603+62+11)=6.44 x 10^-5^).

We performed Mendelian randomisation analyses. As instrument variables, we selected SNPs in CYP3A cluster showing strong associations with CYP3A5 expression in lung (exposure), with a F-statistic above 10 and a p-value under 0.001 (0.05/50 tissues), then removed SNPs in pair with a correlation above *r*^2^ >0.8, estimated on Europeans from GTEx using `ieugwasr` package in R [59], leaving 8 SNPs for analyses. Furthermore, we used the scale function on continuous traits and gene expression to estimate the change of 1 standard deviation (SD) of the phenotypes for 1 SD of the gene expression. As outcomes, we used the 6 phenotypes for which SNPs under selection showed associations for both phenotype (p-value<6.44 x 10^-5^) and CYP3A5 expression (p-value<0.001). Mendelian randomisation analyses were performed using `MendelianRandomization` package in R [60]. We performed Inverse Variance Weighted (IVW) as the main statistical test, and we performed MR-Egger to detect and correct for directional pleiotropy: we report MR-Egger results if the intercept was significant. Lastly, we performed weighted median test as a sensitivity test.

## 7 Author contributions statement

ARSH, IG and JP performed analyses. JCG, RP and IG pre-processed the data. ARSH, IG. and JH wrote the paper, revised by JCG, JP and RP. JGH initiated and supervised the project.

