## Supplementary file for "Signatures of co-evolution and co-regulation in the CYP3A and CYP4F genes in humans"

#### 9 Supplemental Figures

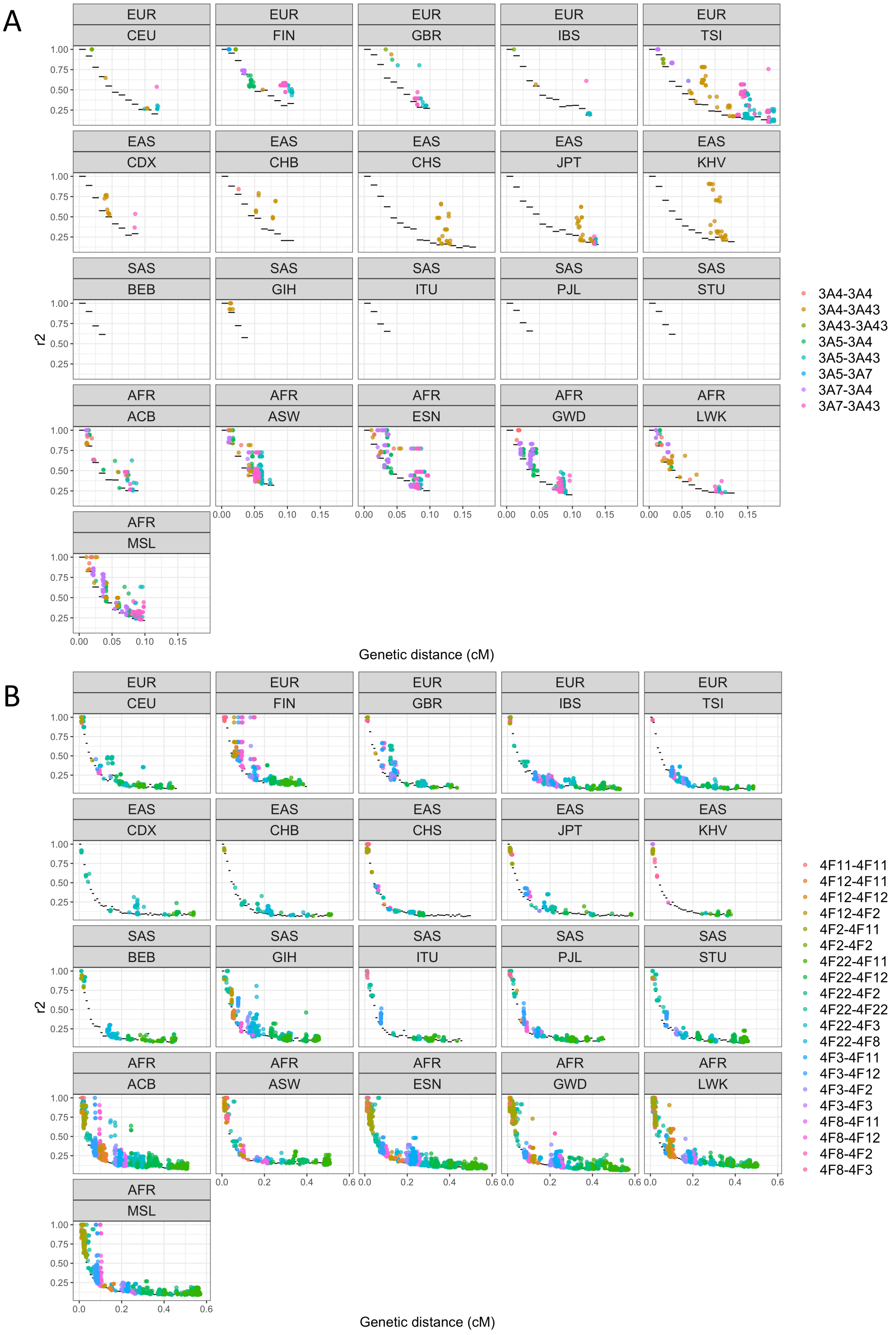


Figure S1: $r^2$ values between each pairs of SNPs in the A) CYP3A and B) CYP4F cluster for each 1000G population, except YRI (AFR). The genetic distance between the SNPs is in centimorgan (cM). Only $r^2$ values over the empirical threshold are shown. The empirical distribution is shown with black horizontal lines. Dots are colored according to which genes are involved in the pair.

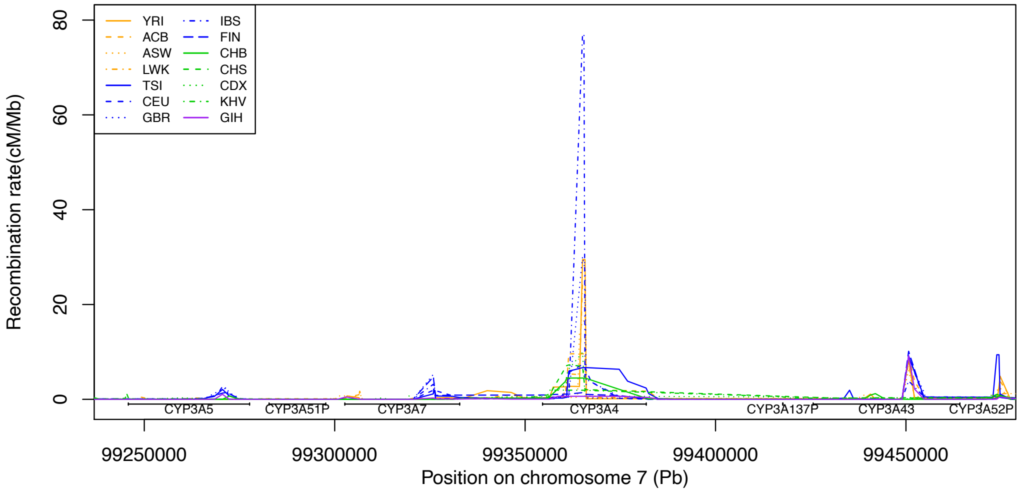


Figure S2: Recombination map in the CYP3A gene cluster. Each line, with a different line pattern, represents a population and is colored according to the super-population. Each gene and pseudogene are shown below the plot with horizontal line.

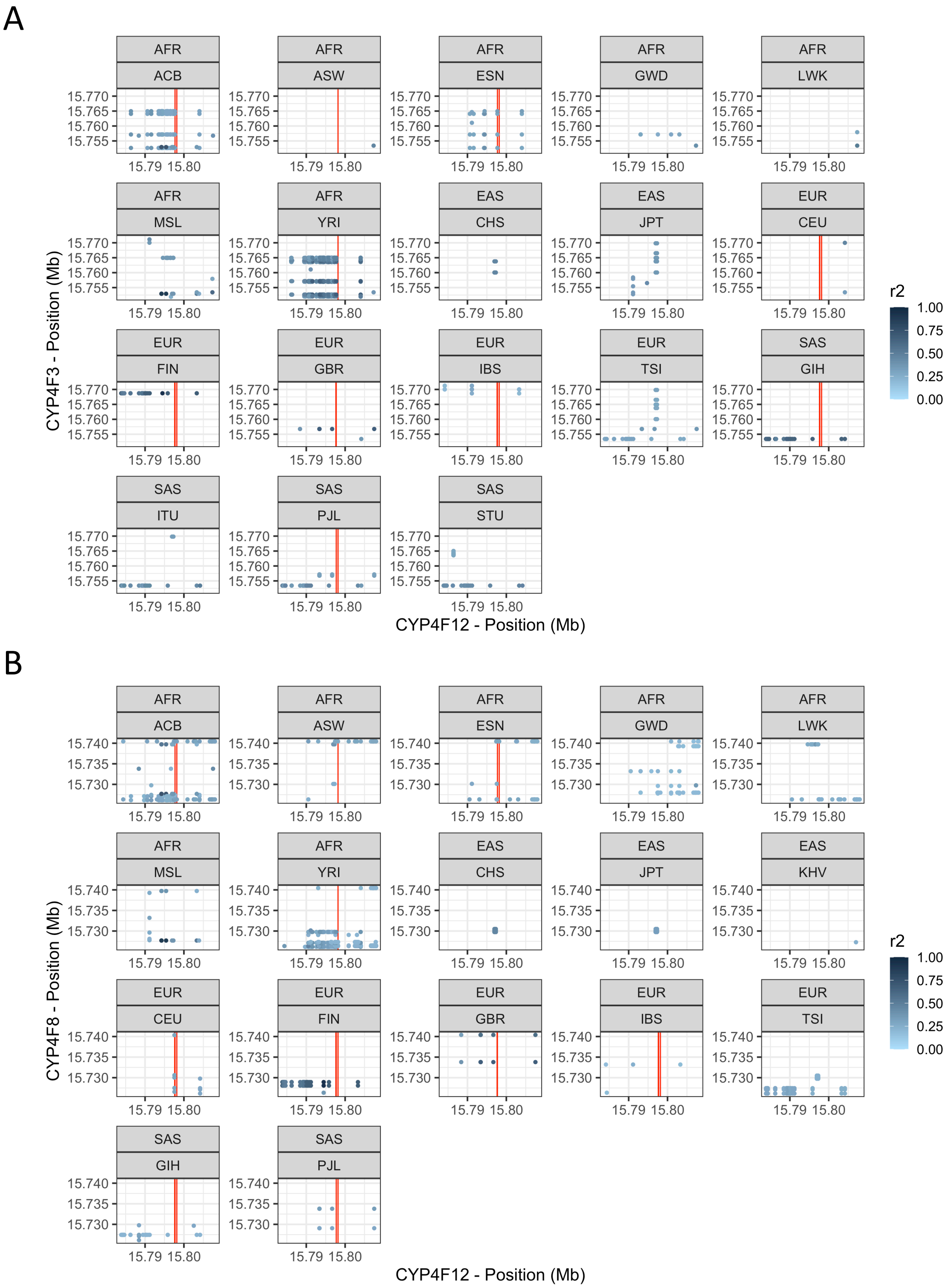


Figure S3: Coordinates of each SNP that is in a pair of SNPs with $r^2$ values in the extremes of the empirical distribution for each subpopulation of 1000G. The displayed SNPs pairs have one SNP in CYP4F12 and the other is in A) CYP4F3 and in B) CYP4F8. We took $r^2$ values from the previous analysis and filtered to keep only values where one SNP was located in CYP4F12. The graph is generated using the *ggplot2* library in R. The physical coordinates of each significant Beta signal, identified in the balancing selection analysis, are shown by the vertical red lines, which were created using *geom_vline*. Points were colored according to their respective $r^2$ values with *scale_color_gradient*.

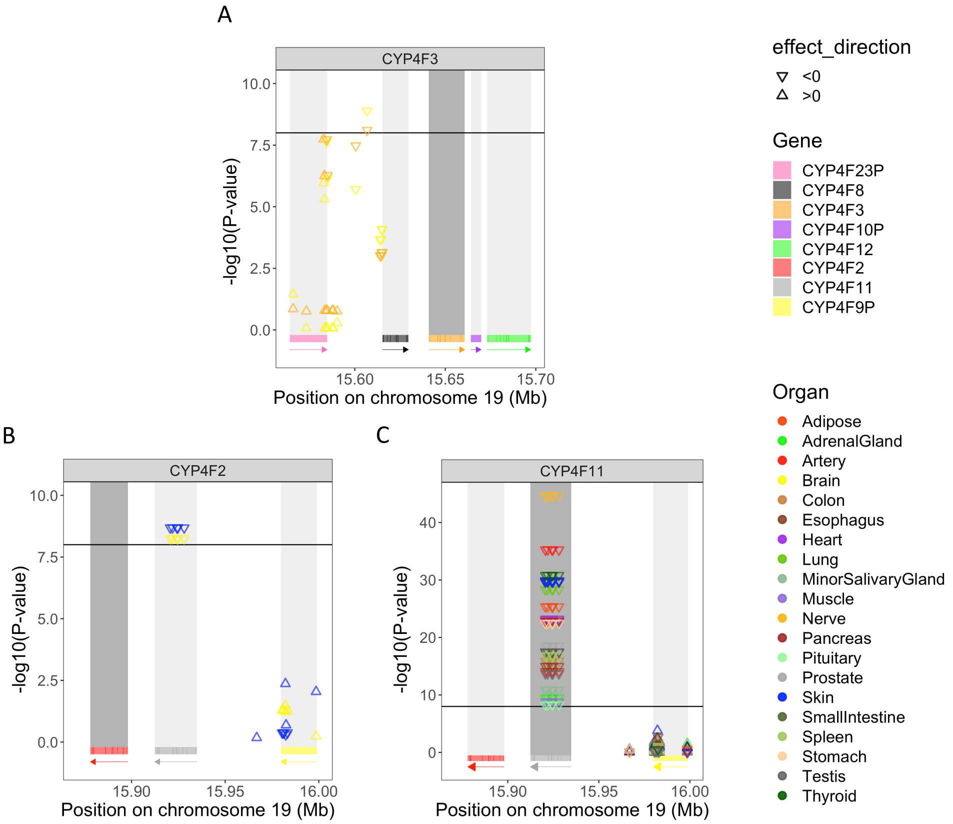


Figure S4: P-values associated with SNPs under positive selection ($|iHS| \geq 2$) explaining variation of gene expression of A) CYP4F3 B) CYP4F2 and C) CYP4F11. The tested gene is shown in dark gray and the effect size is represented either by a triangle standing on its base or a triangle standing on its point. The threshold, set to $10^{-8}$, is represented by the horizontal black line, meaning that a $-log10(p-value)$>8 is a significant eQTL.

| Variant identifier | iHS | Population | Super-population | eQTL |
|---|---|---|---|---|
| rs74459786 | 2.00062 | JPT | EAS | CYP4F12 |
| | 2.09063 | STU | SAS | |
| rs62115147 | -2.09205 | IBS | EUR | CYP4F3 |
| rs2365175 | 2.07818 | TSI | EUR | CYP4F2, CYP4F11 |
| | 2.04181 | KHV | EAS | |
| rs11086013 | 2.11270 | TSI | EUR | CYP4F2, CYP4F11 |
| | 2.01056 | KHV | EAS | |
| rs11881793 | 2.07352 | TSI | EUR | CYP4F2, CYP4F11 |
| | 2.12697 | IBS | EUR | |
| rs3746154 | 2.07395 | TSI | EUR | CYP4F2, CYP4F11 |
| | 2.12768 | IBS | EUR | |
| rs4808413 | 2.06351 | TSI | EUR | CYP4F2, CYP4F11 |
| | 2.17967 | IBS | EUR | |

Table S1: SNPs under positive selection in the CYP4F cluster that are also eQTLs. Each significant SNP is reported with its iHS values ($|iHS| \geq 2$), specific population and RS variant identifier. The gene with differential expression is reported in the eQTL column.

| Variant identifier | $\beta$ score | Population | Super-population |
|---|---|---|---|
| rs644584 | 72.91370 | CEU | EUR |
| | 74.40043 | FIN | |
| | 72.64120 | GBR | |
| | 77.31849 | IBS | |
| | 70.56299 | GIH | SAS |
| rs642322 | 67.17824 | ACB | AFR |
| | 64.12365 | ASW | |
| | 77.53712 | ESN | |
| | 60.42254 | YRI | |
| | 80.95814 | CEU | EUR |
| | 79.46902 | FIN | |
| | 76.20108 | IBS | |
| | 74.54502 | GIH | SAS |
| | 75.44200 | PJL | |
| rs74459786 | 75.65607 | ACB | AFR |
| | 59.94361 | ASW | |
| | 96.13950 | ESN | |
| | 107.11148 | GWD | |
| | 89.83249 | LWK | |
| | 100.15158 | MSL | |
| | 107.11148 | GWD | |
| | 85.28441 | YRI | |
| | 111.07817 | CEU | EUR |
| | 95.36774 | FIN | |
| | 115.01944 | GBR | |
| | 97.56379 | IBS | |
| | 106.51889 | CDX | EAS |
| | 83.53252 | CHS | |
| | 76.40631 | KHV | |
| | 83.65641 | BEB | SAS |
| | 74.91468 | ITU | |
| | 114.13019 | GIH | |
| | 78.72150 | PJL | |
| rs75814017 | 70.44644 | ACB | AFR |
| | 88.88523 | ESN | |
| | 100.74132 | GWD | |
| | 81.90479 | LWK | |
| | 94.13457 | MSL | |
| | 76.91697 | YRI | |
| | 103.15354 | CEU | EUR |
| | 95.98925 | FIN | |
| | 115.75484 | GBR | |
| | 95.16955 | IBS | |
| | 95.82953 | TSI | |
| | 101.29256 | CDX | EAS |
| | 75.79028 | CHS | |
| | 77.93165 | BEB | SAS |
| | 73.87701 | PJL | |

**Table S2 continued from previous page**

| Variant identifier | $\beta$ score | Population | Super-population |
|---|---|---|---|
| rs73000014 | 69.60533 | ESN | AFR |
| | 74.28322 | GWD | |
| | 84.13158 | CEU | EUR |
| | 71.43819 | FIN | |
| | 78.00721 | GBR | |
| | 80.80257 | IBS | |
| | 80.54700 | TSI | |
| | 83.32819 | CDX | EAS |
| | 94.87801 | GIH | |
| rs16980720 | 60.32447 | ACB | AFR |
| | 63.67802 | ESN | |
| | 77.61884 | CEU | EUR |
| | 79.43404 | FIN | |
| | 75.75824 | GBR | |
| | 82.68951 | IBS | |
| | 75.28558 | GIH | SAS |
| | 73.43462 | PJL | |

Table S2: SNPs under balancing selection in the CYP4F cluster that are also eQTLs of CYP4F12. Each significant SNP is reported with its $\beta$ values, specific population and RS variant identifier.

| Description | Code field | Processing |
|---|---|---|
| **Continuous phenotypes suggested by UKb** | | |
| Length of working week for main job | 767 | |
| Frequency of travelling from home to job workplace | 777 | |
| Age completed full time education | 845 | |
| Cooked vegetable intake | 1289 | |
| Salad / raw vegetable intake | 1299 | |
| Fresh fruit intake | 1309 | |
| Dried fruit intake | 1319 | |
| Bread intake | 1438 | |
| Cereal intake | 1458 | |
| Tea intake | 1488 | |
| Coffee intake | 1498 | |
| Water intake | 1528 | |
| Age started wearing glasses or contact lenses | 2217 | |
| Age high blood pressure diagnosed | 2966 | |
| Age diabetes diagnosed | 2976 | |
| Age angina diagnosed | 3627 | |
| Age hay fever, rhinitis or eczema diagnosed | 3761 | |
| Age asthma diagnosed | 3786 | |
| Age heart attack diagnosed | 3894 | |
| Age emphysema/chronic bronchitis diagnosed | 3992 | |
| Age deep-vein thrombosis (DVT, blood clot in leg) diagnosed | 4012 | |
| Age pulmonary embolism (blood clot in lung) diagnosed | 4022 | |
| Age stroke diagnosed | 4056 | |
| Longest period of depression | 4609 | |

**Table S3 continued from previous page**

| Description | Code field | Processing |
|---|---|---|
| Number of depression episodes | 4620 | |
| Age glaucoma diagnosed | 4689 | |
| Age cataract diagnosed | 4700 | |
| Longest period of unenthusiasm / disinterest | 5375 | |
| Number of unenthusiastic/disinterested episodes | 5386 | |
| Age when loss of vision due to injury or trauma diagnosed | 5430 | |
| Age when diabetes-related eye disease diagnosed | 5901 | |
| Age macular degeneration diagnosed | 5923 | |
| Age other serious eye condition diagnosed | 5945 | |
| Hand grip strength (left) | 46 | |
| Hand grip strength (right) | 47 | |
| Waist circumference | 48 | |
| Hip circumference | 49 | |
| Standing height | 50 | |
| Heel bone ultrasound T-score, manual entry | 77 | |
| Heel bone mineral density (BMD) T-score, automated | 78 | |
| Heel bone mineral density (BMD) T-score, automated (left) | 4106 | |
| Heel bone mineral density (BMD) T-score, automated (right) | 4125 | |
| Heel bone mineral density (BMD) T-score, manual entry (left) | 4138 | |
| Heel bone mineral density (BMD) T-score, manual entry (right) | 4143 | |
| Pulse rate | 4194 | |
| Sitting height | 20015 | |
| Fluid intelligence score | 20016 | |
| Birth weight | 20022 | |
| Mean time to correctly identify matches | 20023 | |
| Cascot confidence score | 20121 | |
| Body mass index (BMI) | 21001 | |
| Weight | 21002 | |
| Body fat percentage | 23099 | |
| Whole body fat mass | 23100 | |
| Whole body fat-free mass | 23101 | |
| Whole body water mass | 23102 | |
| Basal metabolic rate | 23105 | |
| Impedance of whole body | 23106 | |
| Impedance of leg (right) | 23107 | |
| Impedance of leg (left) | 23108 | |
| Impedance of arm (right) | 23109 | |
| Impedance of arm (left) | 23110 | |
| Leg fat percentage (right) | 23111 | |
| Leg fat mass (right) | 23112 | |
| Leg fat-free mass (right) | 23113 | |
| Leg predicted mass (right) | 23114 | |
| Leg fat percentage (left) | 23115 | |
| Leg fat mass (left) | 23116 | |
| Leg fat-free mass (left) | 23117 | |
| Leg predicted mass (left) | 23118 | |

**Table S3 continued from previous page**

| Description | Code field | Processing |
|---|---|---|
| Arm fat percentage (right) | 23119 | |
| Arm fat mass (right) | 23120 | |
| Arm fat-free mass (right) | 23121 | |
| Arm predicted mass (right) | 23122 | |
| Arm fat percentage (left) | 23123 | |
| Arm fat mass (left) | 23124 | |
| Arm fat-free mass (left) | 23125 | |
| Arm predicted mass (left) | 23126 | |
| Trunk fat percentage | 23127 | |
| Trunk fat mass | 23128 | |
| Trunk fat-free mass | 23129 | |
| Trunk predicted mass | 23130 | |
| Systolic blood pressure, manual reading | 93 | Values based on the mean of the instance at the first visit |
| Diastolic blood pressure, manual reading | 94 | Values based on the mean of the instance at the first visit |
| Pulse rate (during blood-pressure measurement) | 95 | Values based on the mean of the instance at the first visit |
| Pulse rate, automated reading | 102 | Values based on the mean of the instance at the first visit |
| Forced vital capacity (FVC) | 3062 | Values based on the mean of the instance at the first visit |
| Forced expiratory volume in 1-second (FEV1) | 3063 | Values based on the mean of the instance at the first visit |
| Peak expiratory flow (PEF) | 3064 | Values based on the mean of the instance at the first visit |
| Diastolic blood pressure, automated reading | 4079 | Values based on the mean of the instance at the first visit |
| Systolic blood pressure, automated reading | 4080 | Values based on the mean of the instance at the first visit |
| **Blood cells** | | |
| White blood cell (leukocyte) count | 30000 | |
| Red blood cell (erythrocyte) count | 30010 | |
| Platelet count | 30080 | |
| Lymphocyte count | 30120 | |
| Monocyte count | 30130 | |
| Neutrophill count | 30140 | |
| Eosinophill count | 30150 | |
| Basophill count | 30160 | |
| Nucleated red blood cell count | 30170 | |
| Reticulocyte count | 30250 | |
| High light scatter reticulocyte count | 30300 | |

Table S3: Continuous phenotypes of the UKb

### 10 Supplementary text

#### 10.1 Pre-processing of GTEx genetic data

Starting from the imputed genotyping dataset, we kept bi-allelic SNPs and removed positions with more than 5% missing genotypes, leaving 100,986 SNPs which were used to perform a PCA using flashPCA2 [61]. To retain the non-admixed individuals of European descent, we reduced the dimensionality of the top 10 PCs using the R package `UMAP` [62] (default parameters) to obtain a two dimensional representation of the genetic information contained within those PCs. We identified the largest homogeneous group (self-reported "white") and excluded outlier groups, used only these individuals for the rest of the analyses. We then reran a PCA on this group. We did all subsequent analyses with these 699 individuals. Next we separated each tissue, then removed tissues with fewer than 50 samples, leaving samples from 50 different tissues. We removed in our analyses genes that had fewer than 6 reads in at least 20% of the samples (as recommended by GTEx). We then normalized expression data using limma (TMM normalization) [63] and voom [64]. We calculated PEER factors [65] on the normalized expressions. The suggested number of PEER factors for the GTEx tissues is 15 for $N < 150$, 30 for $150 \leq N < 250$, 45 for $250 \leq N < 350$, and 60 for $N \geq 350$ [66].

#### 10.2 Additional analyses on phenotypes

In CYP4F cluster, no SNP under selection was found associated with one of the selected phenotypes. It is possible that the UKb, and more specifically the white British population, is not the appropriate population in which to investigate these selection signals. However, other SNPs in these loci, not found to be under selection in our previous analyses, are associated with forced expiratory volume in 1 second (FEV1) and forced vital capacity (FVC) in the gene *CYP4F2*, and eosinophil count in *CYP4F23P*.

In the *CYP3A* cluster, we identified significant eQTL signals for genes outside the *CYP3A* family, such as *GS1-259H13.2* (ENSG00000244219.6), *ARPC1B* (ENSG00000130429.12) and *ZKSCAN5* (ENSG00000196652.11), located between *CYP3A7* and *CYP3A4* genes and at the 3' end of *CYP3A43* (Figure 6A. These eQTLs are also associated with anthropometric traits, such as fat and height. None of the other SNPs under selection were found to be associated with the expression levels of these genes, meaning that the associations found could be associated with other members of the *CYP3A* family.

In Mendelian randomisation analyses, there were no causal relationship detected between CYP3A5 expression with neither PID ($p_{IVW}$=0.17), pulse rate ($p_{IVW}$=0.04, $p_{Egger}$=0.72,$p_{Intercept}$=0.02), erythrocyte count ($p_{IVW}$=0.91) and platelet count ($p_{IVW}$=0.11). For pulse rate, the strongest signals were located at the 3' end of CYP3A4, suggesting that this outcome could be associated with expression of another gene in the cluster and/or in another tissue, for which we did not have statistical power to detect appropriate eQTL instruments for Mendelian randomisation.
